# Reanalysis of Alzheimer’s Brain Sequencing Data Reveals Absence of Purported HHV6A and HHV7

**DOI:** 10.1101/830695

**Authors:** Samuel D. Chorlton

## Abstract

Readhead et al. recently reported in *Neuron* the detection and association of Human Herpesviruses 6A (HHV6A) and 7 (HHV7) with Alzheimer’s disease by shotgun sequencing. I was skeptical of the specificity of their modified Viromescan bioinformatics method and subsequent analysis for numerous reasons. In their supplementary data, they report the detection of Variola virus, the etiological agent of the eradicated disease smallpox, in 97.5% of their Mount Sinai Brain Bank dataset. I reanlyzed Readhead et al’s data using highly sensitive and specific alternative methods and find no HHV7 reads in their samples; HHV6A reads were found in only 2 out of their top 15 samples sorted by reported HHV6A abundance. Finally, I recreate Readhead et al’s modified Viromescan method and identify reasons for its low specificity.

## Introduction

I read with great interest the paper from Readhead et al. published in *Neuron* suggesting detection of Human Herpesviruses 6A (HHV6A) and 7 (HHV7) in human brains via shotgun sequencing, along with their association with Alzheimer’s disease (Readhead et al., 2018). This paper has garnered much media attention and spurred a significant amount of research, with 144 citations at the time of writing. Recent literature has supported a role for herpesviruses in the pathogenesis of Alzheimer’s disease, particularly Herpes Simplex Virus 1, as reviewed by Itzchaki (Itzhaki, 2018).

Upon careful reading of the Readhead et al. paper, I was skeptical of their methods for numerous reasons: 1) reporting of uncorrected p-values for multiple testing throughout their paper (eg. figures 2 and 4 in their paper compared with supplementary data); 2) a discrepancy between the reported number of tested viruses in their publication (n=515) and the number of “viruses” reported in their Synapse public data (n=499), many of which are in fact viral segments that collapse into 1 virus (eg. each Influenza virus had 8 genomic segments tested separately) and could have led to multiple testing; 3) failure to correct for multiple testing when testing multiple contrasts (eg viral expression in Definite AD vs Control and Likely AD vs Control were counted as unrelated tests); 4) an extremely low number of putative HHV6A/HHV7 viral reads per sample (eg. in the MAYO_TCX dataset, the maximum number of HHV7 reads in any sample was 5 out of 58 million); and 5) an unusually liberal false discovery rate (0.25) for vQTL analysis, paired with an unsystematic interpretation of vQTL results (despite the use of gene set analysis later in their paper).

After multiple attempts to obtain additional source code and data (personal communication, Feb 6 and Mar 19 2019), I set out to evaluate the presence of human herpesviruses in the data of Readhead et al. using alternative *in silico* analyses and determine the reasons for likely false positive results.

## Methods and Results

Based on supplementary table 2 from Readhead et al, I calculated viral prevalence based on their modified Viromescan (Rampelli et al., 2016). In the main MSBB dataset, HHV6A and HHV7 were detected in 26.8% and 26.3% of samples, respectively, with the authors’ arbitrary cutoff of 2 reads per sample. In comparison, their reported prevalence of Hepatitis C virus in brain tissue was 100% (n=602/602) for the Mount Sinai Brain Brank (MSBB) samples, and 97.5% for Variola virus, the etiological agent of smallpox. Duvenhage virus, which causes a rapid rabies-like death for which I can only find 3 known cases in the literature, was found in 11.2% (n=55/489) of the Memory and Aging Project and Religious Orders Study samples (Paweska et al., 2006; Thiel et al., 2009; Tignor et al., 1977).

Given the implausibly high prevalence of these viruses detected using the modified Viromescan, I next sought to determine if there were any true HHV6A or HH7 reads in the brain samples analyzed by Readhead et al. Source code and data is available at Github: https://github.com/schorlton/Readhead_commentary. Ten samples from each of MSBB_WES (syn7541077), MSBB_RNA (syn8612191) and MAYO_TCX (syn8612203) datasets with the greatest number of reported HHV6A (n=5/dataset) and HHV7 reads (n=5/dataset) were selected for further analysis (n=30 total). Raw reads were preprocessed with fastp, then taxonomically categorized using KrakenUniq, a fast yet highly sensitive method based on k-mers (Breitwieser et al., 2018; Chen et al., 2018). KrakenUniq identified a total of 13 HHV6A reads in 2/15 top HHV6A samples (Readhead total: 75 reads), and failed to identify any HH7 reads in the top HHV7 subset (Readhead total: 93 reads in 15 samples).

KrakenUniq was validated for detection of extremely low viral read counts with a large human background. Thirty samples of 50 million human sequencing reads/read pairs were *in silico* simulated to closely match the MSBB_WES (n=10), MSBB_RNA (n=10) and MAYO_TCX (n=10) datasets (Griebel et al., 2012; Huang et al., 2012). Half the samples from each simulated experiment were spiked with a random HHV6A read, and the other half with a random HHV7 read from any of the public strains not included in the KrakenUniq database. KrakenUniq accurately identified the spiked viral read as present in all samples, and no human reads were classified as HHV6A or HHV7.

As KrakenUniq failed to identify HHV6A/HHV7 reads in almost all samples purported to harbour the most by Readhead et al, I attempted to identify how reads were classified as such by them. While I could not exactly recreate their method and results based on the limited detail in their paper, the median absolute difference in HHV6A/HHV7 readcount from their public data was 4 (IQR:2.5-6.5) in the HHV6A subset and 9 (IQR:4-89.5) in the HHV7 subset. Reads mapping to HHV6A or HHV7 were extracted and subjected to complexity analysis: 55/95 (57.9%) HHV6A and 607/637 (95.3%) HHV7 reads were deemed low complexity by Prinseq(Schmieder and Edwards, 2011). All reads were also subjected to a BLAST search against the non-redundant Genbank (nt) database and collapsed to Lowest Common Ancestor of the top hit (Altschul et al., 1990). Similar to KrakenUniq results, 13 reads from the 15 HHV6A samples aligned to HHV6A, and 0 reads aligned to HHV7. Additionally, when the set of 30 simulated samples, without spiked viral reads, were processed with the modified Viromescan method,10/30 samples had 2 or more reads aligning to HHV6A and no samples had reads aligning to HHV7.

## Discussion and Conclusion

Here I demonstrate that the modified Viromescan used by Readhead et al. likely vastly overestimates viral read counts, and in most cases (28/30 top viral readcount samples), probably identifies viral reads when none are present. Simulation showed this alternative method to be sensitive, and the Readhead et al. method to be highly nonspecific. This latter finding is supported by the detection of an eradicated virus in 97.5% of samples, and BLAST search results of the modified Viromescan’s output. Possible reasons for these findings include abundant low complexity sequences combined with Bowtie2 local alignment, requiring just 27/60 matching bases for a successful alignment, and a 2% false negative rate of bmtagger for filtering human reads (Kirill Rotmistrovsky and Richa Agarwala).

Based on these findings, I suggest that most of the findings based on viral presence presented by Readhead et al. are probably inaccurate, and combined with other statistical flaws, likely falsely reject the null hypothesis. Readhead et al. have failed to respond to my concerns. I suggest that this article be read with significant caution.

## Acknowledgements

None.

## Notes

**Conflicts of interest:** None declared.

https://github.com/schorlton/Readhead_commentary

